# Floral volatiles evoke partially similar responses in both florivores and pollinators and are correlated with non-volatile reward chemicals

**DOI:** 10.1101/2023.02.13.528270

**Authors:** Rohit Sasidharan, Robert R. Junker, Elisabeth J. Eilers, Caroline Müller

## Abstract

**Background:** Plants use floral displays to attract mutualists, but simultaneously need to prevent attacks by antagonists. Chemical displays detectable from a distance include attractive or repellent floral volatile organic compounds (FVOCs). Post-landing, visitors perceive contact chemicals including nutrients, but also deterrent or toxic constituents in pollen and nectar, protecting flowers from overexploitation. The composition of FVOCs and pollen chemistry is well known to vary among and within species. However, we lack knowledge about differences and similarities in the detectability of and behavioural responses towards these compounds for insect flower visitor groups of key importance, i.e., mutualistic pollinators versus antagonistic florivores, as well as the correlation between FVOCs and pollen chemodiversity.

**Scope:** We reviewed how FVOCs and non-volatile floral chemical displays, i.e., nutrients and toxins of pollen, vary in composition and how they affect the detection and behaviour of insect flower visitors. Moreover, we used a meta-analytic approach to evaluate the detection of and responses towards FVOCs by pollinators vs. florivores within the same plant genera. Furthermore, we tested whether the chemodiversity of FVOCs as well as nutrients and toxins stored in pollen are correlated and hence informative about each other.

**Key Results:** According to the available data, florivores are more likely to detect FVOCs than pollinators. Common FVOCs such as linalool and methyl salicylate were often reported as attractive to pollinators and repellent towards florivores. A higher number of FVOCs was found to be attractive to both mutualists and antagonists compared to shared repellent compounds. Furthermore, a negative correlation between FVOC richness and the number of pollen toxin classes was revealed, besides a trend towards a positive correlation between pollen protein amount and the number of pollen toxins.

**Conclusions:** Plants face critical trade-offs when producing floral chemicals, as these partly mediate the same information, particularly attraction but also repellence or toxicity, to both mutualists and antagonists. Moreover, chemodiversity of different floral parts is partly correlated and thus highly relevant for investigations of flower-insect interactions. Further research is needed on more different wild and cultivated plant species and mutualistic and antagonistic interaction partners to test for generalisation of these patterns.

## INTRODUCTION

Animal-pollinated plants face a trade-off with respect to their flower traits: they need to attract pollinators for their reproduction but at the same time prevent visits by florivorous organisms. Thus, they use a broad repertoire of various structural and particularly chemical traits that can act as advertisements and/or defences (Junker, 2016). Floral chemicals comprise both floral organic volatile compounds (FVOCs) which allow signal transmission over distances, as well as stored specialised metabolites present in the flowers, namely in the pollen, nectar or petals, which are only perceived on contact (Knudsen et al., 2006, Palmer-Young et al., 2019, Rivest and Forrest, 2020, Schiestl, 2010). Pollinating flower visitors depend on nutritive pollen and nectar and are attracted to these rewards by both chemical and visual cues (Borghi et al., 2017, Junker and Parachnowitsch, 2015). Next to innate preferences, they may associate display cues with floral resources that serve as rewards (Wright and Schiestl, 2009, Essenberg, 2021). These rewards contain proteins, lipids, carbohydrates and micronutrients (Bonvehi and Jorda, 1997, Vaudo et al., 2020) on which pollinators depend to meet their nutrient requirements.

However, these nutritious sources are also explored by antagonists such as florivores. It should be noted that there is generally a continuous spectrum from mutualistic pollinators to antagonistic florivores, since some pollinators may overconsume costly resources, while florivores may be “costly pollinators”. Moreover, in natural environments, pollen limitation may occur (Aizen and Harder, 2007, Knight et al., 2005, Bennett et al., 2020), suggesting competition between plants for pollen vectors such as insects. As pollen is costly to produce and is exploited as a food source by a large number of animals, including many antagonists, plants cannot afford to produce unlimited amounts of palatable pollen. Instead, they need to prevent excessive consumption, for example, by repelling or not attracting florivores and pollen robbers or by deterrent or toxic chemicals. Defence chemicals used against various target animals should simultaneously not deter or poison pollinators (Lucas-Barbosa, 2016). The task of a flower to attract mutualistic flower visitors and simultaneously remain concealed for antagonistic flower visitors or repel them by their volatile chemical displays has been described as the defence/apparency dilemma (Kessler et al., 2013). Both pollinators and florivores pose a strong selection pressure on floral signals in this regard (Whitehead and Peakall, 2009, Theis and Adler, 2012, Schiestl, 2015).

Producing complex floral blends is potentially one strategy to cope with this dilemma (Kessler et al., 2013). Another strategy of plants is to deploy an “optimal defence” throughout their tissues with specialised metabolites acting repellent, deterrent or toxic being most concentrated in organs or tissues of high value for plant fitness (McCall and Fordyce, 2010). These metabolites may protect the flowers against feeding damage by florivorous insects (McCall and Irwin, 2006) but also against excess pollen removal by pollinators as well as against pathogens (Palmer-Young et al., 2019, Junker and Blüthgen, 2010, Rivest and Forrest, 2020).

Floral scent and pollen chemicals can be quite diverse in terms of both biosynthetic origin and chemistry (Knudsen and Gershenzon, 2006). In general, insect-pollinated flowers are highly chemodiverse in their floral scent profiles in comparison to other outcrossing species (Farré-Armengol et al., 2015). Pollen may also be chemodiverse to some extent (Rivest and Forrest, 2020). Chemodiversity can be measured at different levels, with a focus on the number of compounds (= richness) but also by implementing their evenness, as done, for example, in calculations of the Shannon index or other diversity indices, originally developed for biodiversity of species (Allen et al., 2009, Petrén et al., 2023, Wetzel and Whitehead, 2020). Many studies on floral attractiveness and apparency neglect the role of contact chemicals (Raguso, 2008b). Our knowledge on the role of contact chemicals as feeding stimulants or deterrents for the different types of flower visitors is still largely limited to social bees.

In general, odour and taste cues of flowers are closely linked from an evolutionary and biochemical perspective in shared chemical pathways (Borghi et al., 2017). Moreover, volatile cues advertising non-volatile floral rewards may allow for an associative learning of FVOCs to corresponding rewards (Dobson and Bergström, 2000, Wright and Schiestl, 2009, Hauri et al., 2021, Essenberg, 2021), provided that the flower is receptive (Knauer et al., 2021, Knauer and Schiestl, 2015). Flower chemicals may also serve other functions such as photoprotection or structural optimisation of the flower (Essenberg, 2021). FVOC chemodiversity may be related to the chemodiversity of rewards and may also be informative of nutrient content, but this has, to our knowledge, not been explored. Rather, previous studies looked at the role of individual FVOCs as “honest signals” (Burdon et al., 2020, Haber et al., 2021, Knauer and Schiestl, 2015).

Here, we use a combined approach of a summary of relevant literature and two meta-analyses to understand the diversity and complexity of flower chemicals and ascertain the roles of various chemical displays in impacting the detectability and behaviour of pollinators and florivores. We review and summarise existing literature and highlight potential trade-offs between conflicting processes such as pollination and florivory influenced by chemical displays as well as limitations in the availability of studies concerning florivory, which could skew our understanding of the two processes. Within our meta-analyses, we address two main questions: (1) do responses to FVOCs differ between pollinators (mutualists) and florivores (antagonists) across various plant species? (2) Does floral scent chemodiversity relate to chemical pollen (floral reward) traits? Finally, we provide an outlook of the available information and important points to reflect upon while studying the impacts of floral chemical displays on flower visitors.

## VOLATILE DISPLAYS: FLORAL SCENT

### Complexity and functions of FVOCs

FVOCs are a chemically highly diverse group (Knudsen et al., 2006). The complex floral scent generally comprises a mixture of FVOCs emitted at unique ratios and at temporally highly dynamic rates (Farré-Armengol et al., 2020). For example, a plant may adapt the timing of FVOC emission to daytimes with highest pollinator activity and/or lowest florivore activity (Theis et al., 2007). Moreover, the scent of flowers often changes soon after pollination (Bhattacharya and Baldwin, 2012, Proffit et al., 2018, Schiestl and Ayasse, 2001). Thus, even during the life span of an individual, FVOC emission may be remarkably versatile. Also, FVOCs are not only emitted by the plants themselves, but yeasts and other microbes associated with nectar highly contribute to the odour blend, for which dispersal by flower visitors is crucial (Francis et al., 2021, Farré-Armengol et al., 2020, Cullen et al., 2021). Such microorganisms add to the chemodiversity of FVOCs through metabolising FVOCs or altering plant physiology (Farré-Armengol et al., 2016, Helletsgruber et al., 2017).

Compounds emitted from or occurring in flower parts are not limited to one particular organ. Instead, there exists a high degree of convergence in the production of FVOCs and VOCs from vegetative plant tissues (Stevenson, 2020). Furthermore, to attract pollinators, plants partly produce FVOCs that are identical to compounds involved in animal communication. In fact, an 87% overlap of FVOCs and VOCs produced by insects has been described (Schiestl, 2010). Given the existing overlap between floral and insect scents, it is hypothesised that the specialisation of FVOCs occurred through compound classes that were already detectable by insects (Schiestl, 2010) and that both pollinators and florivores provide strong selection pressures on the evolution of FVOC composition (Strauss and Whittall, 2010, Schiestl, 2010).

FVOCs play multifaceted ecological functions (Farré-Armengol et al., 2020, Junker, 2016, Francis et al., 2021, Cullen et al., 2021). Not only pollinators and floral herbivores, but also predators and parasitoids of herbivores are attracted by certain FVOCs, while other FVOCs repel herbivores or activate resistance against antagonists in neighbouring plants (Sasidharan and Venkatesan, 2020, Farré-Armengol et al., 2020). Since most flower visitors do not specialise on only one plant species, the composition of FVOCs shows great interspecific convergence among plants and is therefore not taxonomically characteristic above the genus level (Schiestl and Johnson, 2013, Knudsen et al., 2006, Farré-Armengol et al., 2020). In fact, some FVOCs are emitted by a huge number of different plant species, of which the monoterpenoids linalool and ocimene and the benzenoid benzaldehyde are among the most commonly reported in both ubiquity and predominance (Knudsen et al., 2006, Farré-Armengol et al., 2020).

### Physiological and behavioural responses towards FVOCs by flower-visiting insects

Floral scent must be perceived by insect visitors against the background of other plant volatiles (Raguso, 2008a). In most insect species, odour-sensitive sensillae are located on the antennae and on the maxillary palps. These sensillae are hair-like structures that contain olfactory receptor neurons (ORNs), also called olfactory sensory neurons (OSNs), and non-neuronal support cells, embedded in lymph fluid sheathed by a porous cuticle (Schmidt and Benton, 2020). The ORNs allow detection and distinguishing of individual molecular structures (Bruce and Pickett, 2011). Based on morphological characteristics such as length, width, cuticle thickness and number and size of pores, different types of sensillae exist, e.g. basiconic or trichoid sensillae (Schmidt and Benton, 2020). Basiconic sensillae are mostly responsible to detect food-derived odours, while trichoid sensillae allow for pheromone detection (Grabe and Sachse, 2018). Both types are presumably responsible for detection of FVOCs in a large number of flower-visiting insects, since FVOCs often resemble food scents or mimic pheromones (Raguso, 2008b). The electrical signals produced as a result of stimulation of sensillae, i.e. neuronal excitation, are experimentally often recorded with the help of electrophysiological tools, in particular electroantennographic detection (EAG), which can be coupled to a gas chromatograph (and mass spectrometer) to reveal the identity of the stimulant compound (GC-(MS-)EAD) (Farré-Armengol et al., 2013). Alternatively, other electrophysiological tools such as single sensilla recordings (SSR) may be used (Sokolinskaya et al., 2020).

Behavioural responses of flower visitors to detected FVOCs depend on the insect’s dependency on floral resources (Junker and Blüthgen, 2010), insect order (Farré-Armengol et al., 2020) as well as the chemical class and concentration of the odour (Terry et al., 2007, Junker and Blüthgen, 2010). For example, obligate flower visitors, usually pollinators, were found on average to respond positively to FVOCs, while facultative, usually antagonistic, visitors showed on average negative responses. For facultative visitors, monoterpenes, alcohols, ethers and ketones mainly act repellent, while benzenoids were often found to be attractive. Such behavioural observations were mostly based on trap, olfactometer and bait assays, and only some were revealed from toxicity or food choice tests (Junker and Blüthgen, 2010).

Different orders of pollinating insects show preferences towards plants that differ in the abundance of various chemical classes. Such findings were revealed among others through modularity analysis of a meta-network of pollinators and associated FVOC classes (Kantsa et al., 2019). For example, coleopteran pollinators generally prefer plants that have higher amino acid-derived FVOCs (Farré-Armengol et al., 2020). Aromatic compounds are more commonly reported to attract lepidopteran pollinators than hymenopteran ones (Schiestl, 2010). Megachilid bees were found to have strong positive associations with sesquiterpenes, Apidae with benzenoids and wasps with C6 green-leaf volatiles and some terpenes (Kantsa et al., 2019). The effects of FVOCs on flower visitors were found to be concentration-dependent. For example, benzenoids are only attractive at particular ranges of concentration, whereas terpenes may be attractive at lower but repellent at higher concentrations (Hao et al., 2013). However, the effects of FVOCs towards pollinators versus florivores have rarely been compared directly, but some examples were found, which are presented in the meta-analysis below.

### Pollinator and florivore responses to FVOCs: meta-analysis 1

To explore whether pollinators and florivores differ in their electrophysiological (EAD) and behavioural responses towards FVOCs within the same plant species, we performed a first meta-analysis. We began by collecting publications that contained information on FVOCs, searching in the ISI Web of Science (1945–2020), Google Scholar, Google Web Search and the database SCENTbase (for details see Supplement S1) and eventually narrowed down our search to those publications (in total 55) that reported on EAD and behavioural responses of insects towards FVOCs. These included 149 FVOCs tested for responses of 42 pollinator species and 65 FVOCs tested for responses of 40 florivore species. The insect species were classified as pollinators or florivores based on available literature of their activity. For eight plant genera, detection or behavioural responses of both associated pollinators and florivores were tested towards the plant-specific FVOCs. These plant genera were *Brassica* (Brassicaceae), *Cirsium* and *Helianthus* (Asteraceae), *Cucurbita* (Cucurbitaceae), *Daucus* (Apiaceae), *Dichaea* (Orchidaceae), *Fragaria* (Rosaceae) and *Nicotiana* (Solanaceae). Together for detection and behaviour, 20 studies were found to report on pollinator responses towards FVOCs from these plants, and 19 for florivores, which were added up into the final analysis.

The percentage of EAD-active FVOCs was calculated for pollinators or florivores for each plant genus and responsiveness compared between the two insect groups in total using a chi-square test. For the behavioural data, the percentage of FVOCs emitting an attraction, repellence or no response was individually calculated for each plant genus and insect group. Again, attractiveness vs. repellence of FVOCs was compared in total amongst insect groups using a chi-square test. Binomial tests were used to examine the distribution of shared attractive and repellent FVOCs between pollinators and florivores. We expected to find a higher number of detectable FVOCs by pollinators rather than by florivores, as outcrossing plants may provide diverse and conditionally also alternating mixes of compounds targeted at pollinators, while pollinators may have acquired the ability to detect and to associate those compounds with nutrition faster than florivores (Kessler et al., 2013) Regarding behavioural responses, we expected to find a higher proportion of FVOCs to be attractive to pollinators when compared to florivores, as plants must minimise trade-offs between pollinator attraction and florivore repellence to maintain the fitness benefits of pollination.

Our meta-analysis 1 revealed ten most commonly tested FVOCs, which were tested against pollinators and florivores of >= 3 plant genera, for both detection and behavioural responses (Table 1). These are also amongst the most commonly reported FVOCs across angiosperms (Farré-Armengol et al., 2020, Knudsen and Gershenzon, 2006). In total, FVOCs (including unknown compounds) were tested uniquely (against a specific insect) 220 times for pollinators and 102 times for florivores in detection tests (including unknown compounds) from the selected plant genera. Tested pollinators (belonging to 16 species) responded to 151 of these tests (= 69%), while tested florivores (21 species) responded to 83 of these compounds (= 81%), which corresponds to 62 and 32 FVOCs respectively (Table 2). Thus, florivores were significantly more likely to detect tested FVOCs than pollinators (*χ*^2^ = 5.069, *p* = 0.024). This pattern was found in most genera, except for *Cucurbita* and *Fragaria*, which may either represent different specificity across plant taxa, or point towards a limitation in the availability of studies with pollinators and florivore detection responses.

**Table 1.** Most abundant FVOCs tested for both detection and behaviour in >= 3 plant genera against at least one pollinator and florivore species. Poll = Pollinator, Flor = Florivore. Values represent percentages of the total number (in parentheses) of insect species tested for detection or behaviour. For a given compound an insect is sometimes repeatedly tested (e.g., *Apis mellifera*) but the response was found to be uniform and hence averaged below.

| Compound | Detection |  | Behaviour |  |  |  |  |  |
| --- | --- | --- | --- | --- | --- | --- | --- | --- |
|  |  |  | Attracted (+) |  | Repelled (-) |  | No response (0) |  |
|  | Poll | Flor | Poll | Flor | Poll | Flor | Poll | Flor |
| 2-Phenylethanol | 77.8 (9) | 100 (3) | 66.7(3) | 33.3 (3) | 0 (3) | 33.3 (3) | 33.3 (3) | 33.3 (3) |
| 3-Hexenol | 100 (4) | 100 (1) | 14.3 (7) | 14.3 (7) | 14.3 (7) | 0 (7) | 71.4 (7) | 85.7 (7) |
| Benzaldehyde | 100 (9) | 100 (4) | 33.3 (9) | 27.3 (11) | 0 (9) | 0 (11) | 66.7 (9) | 72.7 (11) |
| Benzyl alcohol | 87.5 (8) | 100 (2) | 28.6 (7) | 20.0 (10) | 0 (7) | 0 (10) | 71.4 (7) | 80.0 (10) |
| Hexanol | 100 (3) | 100 (2) | 0 (2) | 50.0 (2) | 100 (2) | 0 (2) | 0 (2) | 50.0 (2) |
| Limonene | 66.7 (3) | 100 (3) | 50.0 (2) | 50.0 (2) | 0 (2) | 0 (2) | 50.0 (2) | 50.0 (2) |
| Linalool | 90.0 (10) | 100 (2) | 22.2 (9) | 0 (8) | 0 (9) | 12.5 (8) | 77.8 (9) | 87.5 (8) |
| Methyl salicylate | 100 (10) | 100 (4) | 37.5 (8) | 0 (9) | 0 (9) | 11.1 (9) | 62.5 (8) | 88.9 (9) |
| Nonanal | 100 (3) | 100 (1) | 50.0 (2) | 0 (1) | 0 (2) | 0 (1) | 50.0 (2) | 100 (1) |
| Phenylacetaldehyde | 85.7 (7) | 100 (6) | 28.6 (7) | 41.7 (12) | 0 (7) | 0 (12) | 71.43 (7) | 58.3 (12) |

**Table 2.** Differences between responses of pollinators and florivores to tested FVOCs of different plant genera. Percentage of FVOC tests, which were electrophysiologically detected or acted attractive or repellent in behavioural studies (% of *n*) and total number (*n*) of unique FVOC tests with insect species within that category are given. Total represents every unique combination of tested FVOC-insect. Proportions differing significantly (*p* ≤ 0.05, chi-square test) for the total between pollinators and florivores are highlighted in bold. No response was included as a category in tests for attractive/repellent FVOCs but the data is not shown below. NT = not tested. Data from 32 studies in total.

| Plant | Response | Pollinator |  | Florivore |  |
| --- | --- | --- | --- | --- | --- |
|  |  | % of total | total *n* | % of total | total *n* |
| *Brassica* spp. (*B. nap*a, *B. rap*us) | Detected | 91.3 | 23 | 100 | 8 |
|  | Attractive | 52.0 | 25 | 75.0 | 4 |
|  | Repellent | 4.0 |  | 0 |  |
| *Cirsium* spp. (*C. arven*se, *C. japonicum*) | Detected | 70.4 | 81 | 90.9 | 11 |
|  | Attractive | 24.1 | 54 | 9.1 | 66 |
|  | Repellent | 0 |  | 3.0 |  |
| *Cucurbita* spp. (*C. maxima*, *C. pepo*) | Detected | 100 | 2 | 96.6 | 29 |
|  | Attractive | 66.7 | 3 | 30.8 | 65 |
|  | Repellent | 0 |  | 1.5 |  |
| *Daucus carrota* | Detected | 23.1 | 21 | 75.0 | 8 |
|  | Attractive | 100 | 3 | 0 | 0 |
|  | Repellent | 0 |  | 0 |  |
| *Dichaea pendula* | Detected | 0 | 0 | 100 | 1 |
|  | Attractive | 0 | 1 | 100 | 1 |
|  | Repellent | 0 |  | 0 |  |
| *Fragaria* spp. (*F. vesca*, *F. x ananassa*) | Detected | 100 | 13 | 58.3 | 36 |
|  | Attractive | 0 | 4 | 50.0 | 2 |
|  | Repellent | 50.0 |  | 0 |  |
| *Helianthus annuus* | Detected | 68.5 | 54 | 100 | 1 |
|  | Attractive | 100 | 2 | 100 | 1 |
|  | Repellent | 0 |  | 0 |  |
| *Nicotiana attenuata* | Detected | 68.0 | 50 | 100 | 10 |
|  | Attractive | 26.1 | 23 | 19.1 | 21 |
|  | Repellent | 17.4 |  | 28.6 |  |
| Total | Detected | **68.6** | 220 | **81.4** | 102 |
|  | Attractive | 33.0 | 112 | 22.0 | 159 |
|  | Repellent | 8.0 |  | 5.7 |  |

$p = 0.024$ $(\chi^2 = 5.069)$ $p = 0.07$ $(\chi^2 = 5.33)$

The fact that florivores seem to physiologically detect a higher percentage of tested FVOCs than pollinators could be an outcome of stronger pollinator specialisation on floral scent (Salzmann et al., 2006, Farré-Armengol et al., 2020). Pollinators may thus rely on detecting a more limited set of corresponding FVOCs that specially evolved towards attracting them, rather than florivores (Schiestl, 2015). The latter may have randomly adapted to diverse FVOCs. Also, pollinators are often far more mobile than herbivores including florivores (Theis and Adler, 2012, Underwood et al., 2020). Thus, pollinators can scout larger distances to find floral resources and potentially afford to detect a more limited number of FVOCs. In contrast, florivores cannot afford a prolonged foraging time due to their limited mobility, and may thus need to be responsive to a larger number of FVOCs. Lastly, it is important for florivores to sensitively detect toxic compounds produced by plants against them and this may have spilt over into their detection of FVOCs, while some pollinators might be resistant or tolerant to the same compounds (Barlow et al., 2017, Wright et al., 2010).

With regard to behavioural responses, no general differences were observed in the attractiveness of FVOCs towards pollinators in comparison to florivores in the tested plant genera. We found 112 unique tests (as an unrepeated combination of a given FVOC tested against a given insect species) of attractiveness towards pollinators, and 159 tests towards florivores. Most of the tests were performed using choice, olfactometer or proboscis extension reflex (PER) assays, usually comparing preferences for blank control versus an FVOC odour (Table 1). Towards several of the ten most commonly tested FVOCs, relatively more pollinator species seemed to be attracted, while more florivore species were repelled, although this could not be statistically tested as the numbers of tests were low (Table 1). Among these FVOCs, the terpene linalool is reported to act as repellent or deterrent towards florivores in several studies (Junker et al., 2010a, Junker et al., 2011). Interestingly, our analysis revealed that an unrelated but commonly occurring benzenoid, methyl salicylate, also seems to play a similar role in attracting floral pollinators while repelling florivores. While trade-offs between attracting pollinators and repelling florivores are hypothesised to exist (Farré-Armengol et al., 2013), compounds such as linalool and methyl salicylate may offset such trade-offs by causing distinct responses in both groups of flower visitors. One proposed explanation may be that mutualistic pollinators often are obligate flower visitors, while antagonistic flower visitors are often facultative, leading to a co-evolutionary pathway where an FVOC could act attractive to pollinators in general or repellent to florivores in general (Junker and Blüthgen, 2010), with exceptions depending on feeding preferences. Alternatively, such distinct responses to individual floral VOCs may be evident when pollinators and florivores belong to different insect orders (Farré-Armengol et al., 2020, Harrewijn et al., 1994, Schiestl, 2010). In our meta-analysis 1, the majority of florivores tested in the various studies belonged to the Coleoptera and a few studies were performed on florivores belonging to Hymenoptera, Lepidoptera, Orthoptera and Hemiptera, whereas the majority of tested pollinators belonged to the Hymenoptera and Lepidoptera and some to the Diptera. The phylogenetic status and the functional role in relation to floral resources (i.e., pollination vs florivory) may thus be related (Schiestl, 2010). Taken together, it seems that only a few compounds in floral bouquets mediate contrasting responses in some pollinators and florivores. Observations of insect responses are biased towards FVOCs that are more abundantly detected by tools such as GC-MS or GC-flame ionisation detection (FID). However, more tests are needed on “less abundant” FVOCs. Bioassays with manipulated FVOC bouquets using floral extracts and interaction network analyses might help elucidating the role of less abundant FVOCs and the complex networks between plants and flower-visitors (Junker et al., 2010b, Larue et al., 2016, Kantsa et al., 2019).

While few FVOCs cause contrasting responses, significantly more of the tested FVOCs emitted by the selected eight plant genera act attractive rather than repellent on both groups of flower visitors, pollinators and florivores (Table 3). Attractive FVOCs likely evolved primarily to attract pollinators, while repellent FVOCs may have evolved under both pollinator and florivore-imposed selection pressures. Since florivores eavesdrop on floral cues meant to attract pollinators, attractive (rather than repellent) FVOCs could represent a major source of the trade-off faced by flowering plants. However, the low number of studies, particularly for florivores, makes it difficult to determine a clear separation of pollinator-attractive and pollinator/florivore-repellent FVOCs.

**Table 3.**
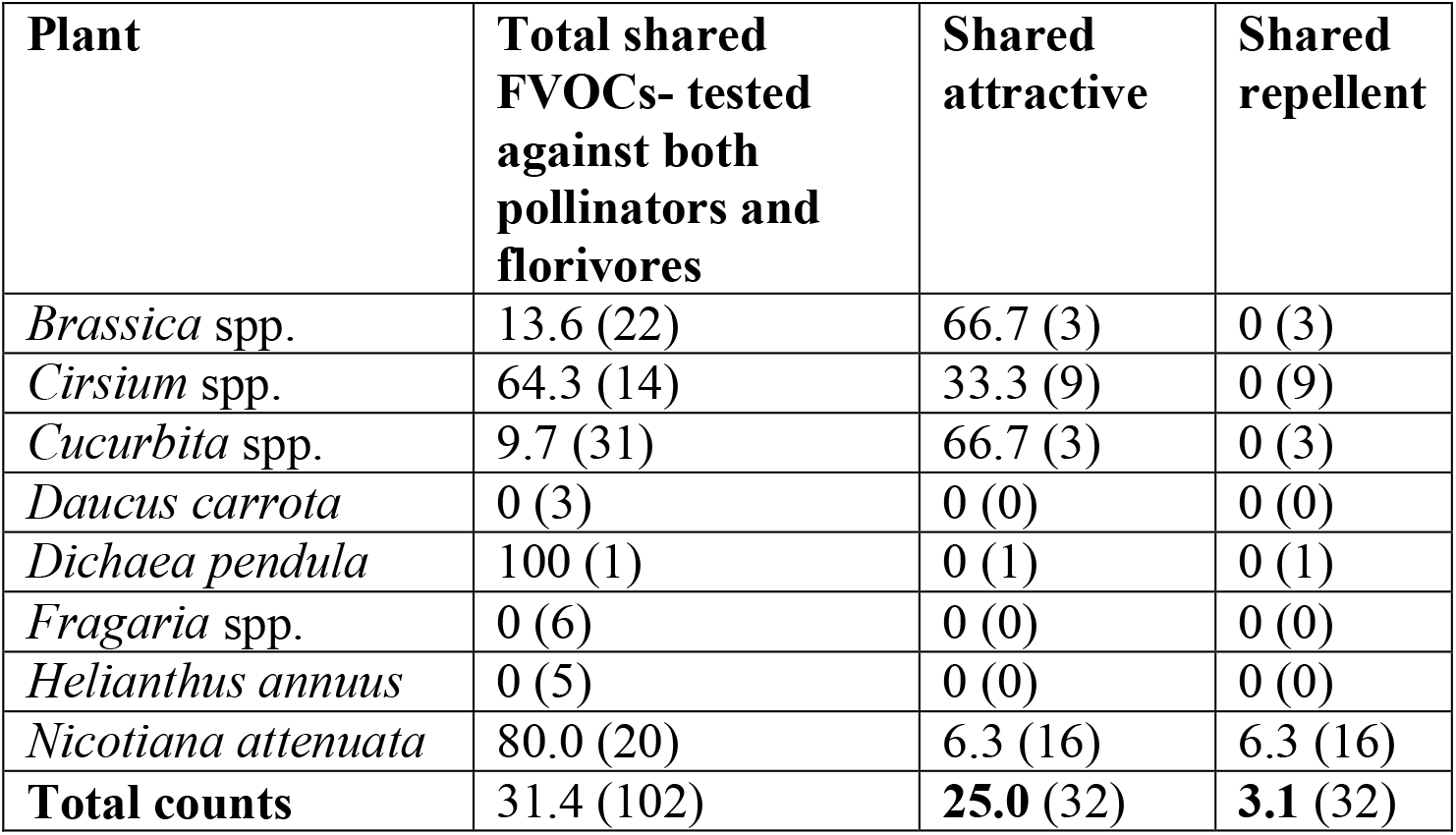
Shared total, attractive and repellent FVOCs of eight plant genera tested for behavioural responses of pollinators and florivores. Values represent percentages of the numbers (in parentheses) of FVOCs tested. Proportions highlighted in bold indicate significant differences (*p* = 0.04, binomial test) in attractive versus repellent shared FVOCs.

| **Plant** | **Total shared FVOCs- tested against both pollinators and florivores** | **Shared attractive** | **Shared repellent** |
| --- | --- | --- | --- |
| *Brassica* spp. | 13.6 (22) | 66.7 (3) | 0 (3) |
| *Cirsium* spp. | 64.3 (14) | 33.3 (9) | 0 (9) |
| *Cucurbita* spp. | 9.7 (31) | 66.7 (3) | 0 (3) |
| *Daucus carrota* | 0 (3) | 0 (0) | 0 (0) |
| *Dichaea pendula* | 100 (1) | 0 (1) | 0 (1) |
| *Fragaria* spp. | 0 (6) | 0 (0) | 0 (0) |
| *Helianthus annuus* | 0 (5) | 0 (0) | 0 (0) |
| *Nicotiana attenuata* | 80.0 (20) | 6.3 (16) | 6.3 (16) |
| **Total counts** | 31.4 (102) | **25.0 (32)** | **3.1 (32)** |

All plant species included in our analysis are outcrossers or at least highly benefit reproductively from outcrossing and are visited by a number of different insect taxa. This pattern may well be representative for various plant-insect relationships, as most outcrossing plants are visited by mutualistic and antagonistic insects of different taxa and degrees of specialisation. Nevertheless, there is a clear need for more studies looking at florivores and their responses to FVOCs in other plant genera. Responses to FVOCs should also be compared in plants that are pollinated by specialists, such as nursery pollination systems (Dötterl et al., 2006) or in plants that make use of mixed-mating and are highly flexible in pollinator attraction.

### Context-dependency of insect responses towards FVOCs

Individual flower visitors can respond differently to FVOCs depending on the context. The composition of available odour receptors and thus the number of FVOCs that are receptor-detected is species-specific. Together with the specificity in neuronal or higher cognitive processes, this can lead to a strong receiver bias (Schiestl and Johnson, 2013). Yet even within the same insect species, the ability to detect a compound and respond behaviourally towards it depends on internal and external conditions, such as food supply and stress. In humans, a high chemosensory plasticity exists, depending for instance on age, disease state and environmental exposure (May and Dus, 2021). Similar levels of plasticity exist in insects. For instance, starvation induces enhanced olfactory detection and transcriptional changes encoding chemosensory proteins and odorant-degrading enzymes in *Spodoptera littoralis* caterpillars (Poivet et al., 2021). In small-scale diverse natural environments and for flower visitors that differ in age, sex, genetic origin and other factors, odour perception is highly subjective and temporally dynamic, even within the same insect species, forming individual “insect odourscapes” in the vicinity of flowers (Conchou et al., 2019). The difficulty of breaking down this complex picture into simple rules is the main limitation of our understanding of the relationship between FVOCs and the different flower visitors. Beyond this natural plasticity in flower chemistry and visitor behaviour, the presence of anthropogenic pollutants such as agrochemicals complicates the relationships, as these can impede the foraging activity of insects (Kessler et al., 2015).

Moreover, single compounds in the mix of emitted FVOCs interact in the odour plume above flowers and insect responses to a whole blend often differ to responses towards individual components (Bruce and Pickett, 2011). Single FVOCs may not be recognised as host cues when perceived outside the context of the entire bouquet. However, FVOCs are often tested individually for detectability in EADs and for behavioural responses. In order to elucidate, which compounds mask or eliminate the attractive effect of other FVOCs, corresponding mixtures have to be investigated. Further studies are required to understand flower-insect interactions under complex environmental settings in order to account for contextual dependency of responses and plasticity of both FVOCs and insect behaviour.

## CONTACT CHEMICAL DISPLAYS: NUTRIENTS AND TOXINS IN POLLEN

### Complexity and functions of pollen chemicals

Plant pollen contains 2.5% to 61% proteins and 2% to 20% lipids (Roulston et al., 2000, Annoscia et al., 2017), while the remaining portion consists mainly of carbohydrates, such as starch (Roulston et al., 2000) and the sugars fructose, glucose and sucrose (Bonvehi and Jorda, 1997). However, the chemical composition of pollen varies greatly depending on plant phylogeny (Roulston and Buchmann, 2000, Vaudo et al., 2020), seasonal appearance and pollination type. For instance, pollen from spring- and autumn-blooming plants contain higher protein proportions compared to pollen from summer-blooming plants (Roulston et al., 2000, Liolios et al., 2015). The pollen protein content is notably higher in pollinator-dependent plants compared to plants with low insect dependence (Ruedenauer et al., 2019). Thus, the protein content of pollen is generally considered as the most important quality characteristic for pollen-feeding insects. However, a correlation between pollen protein content and insect behavioural preferences is not apparent across different species of plants and flower visitors (Roulston et al., 2000). At least for bees, a clear pattern seems to emerge when focussing on the ratio of the main pollen macronutrient components; they generally prefer pollen with an average protein to lipid ratio of 2.5:1 (Vaudo et al., 2016, Vaudo et al., 2020).

Nutrients in pollen do not act solely as rewards for the purpose of positive reinforcement of pollinators, but also serve an important physiological function in the fertilisation process of flowers. Namely, pollen proteins enable pollen tube formation (Roulston et al., 2000) and pollen lipids are required to penetrate the stigma (Wolters-Arts et al., 1998). Thus, pollen nutrients must be considered for their multiple functions in both interactions of plants with flower visitors and endogenous purposes.

In addition to compounds of nutritional value, chemicals in pollen and other flower parts also comprise deterrent or toxic compounds, which belong to the same chemical classes as those produced for defence against herbivores elsewhere in the plant (Stevenson, 2020, Rivest and Forrest, 2020). Most common classes of specialised metabolites in pollen are alkaloids, terpenoids and phenolic compounds, such as flavonoids and phenylpropanoids (Palmer-Young et al., 2019). Some of these may act as toxins when present in sufficiently high concentrations (Palmer-Young et al., 2019, Rivest and Forrest, 2020). Toxins in floral tissue deter florivores and can thus reduce floral damage (Adler et al., 2001, Austel et al., 2021, Irwin and Adler, 2006, Strauss et al., 2004). In many of these cases, such compounds simultaneously deter beneficial pollinators (de Mesquita et al., 2010, Irwin and Adler, 2006, Irwin et al., 2014). However, those toxins are also believed to be important in driving the specialisation of certain bees as pollinators for plants (Rivest and Forrest, 2020).

### Physiological and behavioural responses towards pollen chemicals by flower-visiting insects

For physiological perception of contact chemicals, insect species possess taste organs, so-called gustatory sensillae, mostly on their tarsi, antennae and mouthparts (Bestea et al., 2021, Mitchell et al., 1999). These sensillae are similar in structure to olfactory sensillae, contain gustatory receptor neurons (GRN) and non-neuronal support cells embedded in lymph fluid, but the sheathing cuticle is uniporous (Mitchell et al., 1999, de Brito Sanchez et al., 2007, Bestea et al., 2021). By repeated experiences, flower visitors can learn, to a species-specific degree, to associate visual and volatile cues to encountered contact chemicals (Reinhard et al., 2004), such as rewards, deterrents and toxins. However, particularly little is known about how and to what extent non-volatile floral compounds such as pollen nutrients can be detected and act as feeding stimulants or deterrents to different flower visitor species (Muth et al., 2016). Honeybees are the most studied flower visitors for their sense of taste. Responses to sugars, salts, amino acids (Jung et al., 2015) and fatty acids were found in honeybees (Ruedenauer et al., 2021), while the antennal gustatory sensillae are less likely to respond to bitter substances (de Brito Sanchez et al., 2005, de Brito Sanchez et al., 2007). In fact, several pollinators may lack gustatory receptors that perceive a bitter taste, but still show an avoidance response to such tastes (Ayestaran et al., 2010, Guiraud et al., 2018), which may be due to the inhibition of sweet-responding gustatory receptors (Bestea et al., 2021).

With regard to behavioural responses, preferences of flower visitors are generally investigated towards various pollen components. Preferences for protein contents in pollen differ seasonally and depend on the insect species (DeGrandi-Hoffman et al., 2021, Liolios et al., 2015). For instance, honeybees prefer protein-rich pollen late in the season (Quinlan et al., 2021). Pollen amino acid composition may affect the foraging behaviour of honeybees (Cook et al., 2003) but seemingly not of common bumblebee species (Kriesell et al., 2017). Instead, bumblebees use pollen lipid content as a cue to assess overall diet quality and maximise their fitness (Ruedenauer et al., 2020). Furthermore, pollen lipid constituents, such as the free fatty acids oleic acid, alpha-linolenic acid or linoleic acid, affect foraging behaviour by enhancing the visual learning ability in honeybees and bumblebees (Muth et al., 2018, Zarchin et al., 2017). Moreover, pollen lipids can improve the resilience of bees to pathogen and parasite infestations (Crone and Grozinger, 2021, Annoscia et al., 2017, Schmehl et al., 2014, Barraud et al., 2020, Dolezal and Toth, 2018, Tosi et al., 2017). Contrarily, augmented lipid ingestion increases bumblebee mortality (Ruedenauer et al., 2020). In addition to pollen macronutrients, pollen micronutrients can play a role for flower visitor preferences. For instance, sodium and potassium in pollen influence the preferences of honeybees (Khan et al., 2021). Fewer studies exist on the effects of nutritional components on florivores. Amino acids such as L-alanine and L-serine were found to evoke strong electrophysiological and behavioural responses in the western corn rootworm (Hollister and Mullin, 1998). Sugars evoke an additive positive behavioural response in this florivore species (Kim and Mullin, 2007). However, the feeding behaviour and responses to these available nutrients may be altered by the presence of toxic specialised metabolites, as suggested for the facultative florivore *Coleomegilla maculata* (Lundgren and Wiedenmann, 2004).

Toxins in floral tissues may also be of value to flower visitors for defence or communication purposes. For example, pollen toxins can repel or deter parasites and bees protect their pollen stores by these toxins (Spear et al., 2016). A contribution of the pollen diet for bumblebees by “low quality” (Nicolson and Human, 2013) and flavonoid-rich (Fatrcová-Šramková et al., 2016) sunflower pollen reduces the infection prevalence with a protozoan pathogen (Giacomini et al., 2021). Likewise, the consumption of nectar alkaloids can reduce the protozoan pathogen load in bumblebees (Manson et al., 2010). Although toxins in pollen are believed to be largely targeted against florivores (Adler et al., 2001, Austel et al., 2021, Irwin and Adler, 2006, Strauss et al., 2004), pollinators may nonetheless also be affected by pollen and nectar toxins at certain concentrations, leading to deterrence, impaired development or even death (de Mesquita et al., 2010, Irwin and Adler, 2006, Irwin et al., 2014, Stevenson, 2020). Thus, a similar trade-off as found in FVOCs may exist in pollen traits, namely, the need to protect nutrients serving as rewards for pollinators with toxins to deter florivores and excessive pollen consumers.

Potentially deleterious effects of suboptimal pollen nutrient composition may be overcome by mixing low and high quality resources (Bukovinszky et al., 2017). Furthermore, most flower visitors mix pollen with nectar, leading to nutritional complementarity (Blüthgen and Klein, 2011). Nectar is an important source of energy and should thus be considered in this context (Nicolson, 2011). Moreover, individual harmful compounds such as toxins can be diluted by mixing pollen sources (Eckhardt et al., 2014). However, this strategy of dietary mixing may be used by both groups of flower visitors, pollinators and florivores; thus, the trade-off may be difficult to resolve for plants.

### Relationships between FVOCs and pollen chemicals: meta-analysis 2

FVOCs and pollen chemicals are mostly studied independently, while little is known about potential relationships between FVOCs and other floral chemical traits. Some studies suggest that floral signals are often informative of reward trait quality or composition (Dobson and Bergström, 2000, Hauri et al., 2021, Zu et al., 2022, Essenberg, 2021). Moreover, FVOC production and that of stored toxins could be biosynthetically linked (Dudareva et al., 2013, Pichersky et al., 2006, Twaij and Hasan, 2022). Thus, we expected that the chemodiversity of FVOCs and pollen may be correlated. Flower visitors often respond to blends of FVOCs (Bruce and Pickett, 2011, Kárpáti et al., 2013) and if FVOC chemodiversity is related to pollen chemodiversity, FVOC diversity could inform pollinators or florivores about the nutrient status or toxicity of pollen resources, allowing them to determine the quality at a distance. Thus, we performed a second meta-analysis, using the same literature survey as mentioned above, to test whether FVOC chemodiversity relates to pollen traits, namely nutrient content and chemodiversity of toxins known from pollen. Studies were selected based on the joint availability of data for plant species on the FVOC profile (as a measure of FVOC chemodiversity), their pollen protein content (as a measure of resource quality), and toxins found in pollen. As a measure for chemodiversity of FVOCs, we used the compound richness (i.e., number of FVOCs) and the Shannon diversity (*H*_s_ = -Σ *p*_i_ * ln *p*_i_, where *p* is the relative abundance of an FVOC i), calculated based on the FVOC data from various sources, and averaged for the different values for the individual plant species. Pollen toxins were grouped into their respective chemical classes (e.g., alkaloids, terpenoids etc.). Correlations between the different traits were tested using Pearson’s or Spearman’s rank correlation analyses, depending on whether the data was parametric or not (Supplement S1).

For 49 plant species data on FVOC composition and pollen protein amounts were available. Neither the FVOC richness (Pearson correlation, *r*^*2*^ = 0.017, *p* = 0.36), nor the FVOC Shannon diversity index (Spearman correlation, *ρ* = −0.073, *p* = 0.62) were correlated to the amount of protein in pollen (weight of protein per dry weight (d.w.) of pollen in mg g^-1^) across these studies (*n* = 49, Fig. 1A and Fig. 1B). Sixteen studies were found to report on FVOCs and pollen toxins. Across these studies, compound richness of FVOCs was negatively correlated with the number of stored toxins in pollen within the same plant species (Spearman correlation, *ρ* = −0.587, *p* = 0.02). However, the Shannon indices of the FVOCs were not correlated with the number of stored toxins (Spearman correlation, *ρ* = −0.18, *p* = 0.50; Fig. 1C and Fig. 1D). Similarly, the compound richness of FVOCs was found to be negatively correlated with the number of pollen toxin classes (Spearman correlation, *ρ* = −0.625, *p* = 0.01), while the Shannon index was not (Spearman correlation, *ρ* = −0.377, *p* = 0.15; Fig. 1E and Fig. 1F). The negative correlation between the richness of FVOCs and pollen toxins may be informative, such that more chemodiverse flowers are less toxic and therefore attractive to a wider array of flower visitor species. Alternatively, the negative correlation may be a consequence of metabolic or biosynthetic trade-offs between floral volatiles and specialised metabolites stored in pollen. However, costs may be minimised if compounds are biosynthetically related or chemically similar (Wetzel and Whitehead, 2020, Whitehead and Peakall, 2009). Whether plants divert more chemodiversity into their FVOCs or stored toxins may depend on the individual needs of the plant within its ecological niche and the plethora of visitors with which it must interact. For example, *Prunus dulcis* (almond) has a high diversity in pollen toxins and a low reported FVOC diversity. Almond plants depend on generalised bees for pollination early in the season, when frost damage poses a danger (Henselek et al., 2018). Because almond has many bee pollinators, it may employ more defence chemicals that enhance its resistance towards florivory, but fewer FVOCs that are adequate to attract its generalised bee pollinators to effectively pollinate the flowers quickly (Henselek et al., 2018).

**Figure 1.**
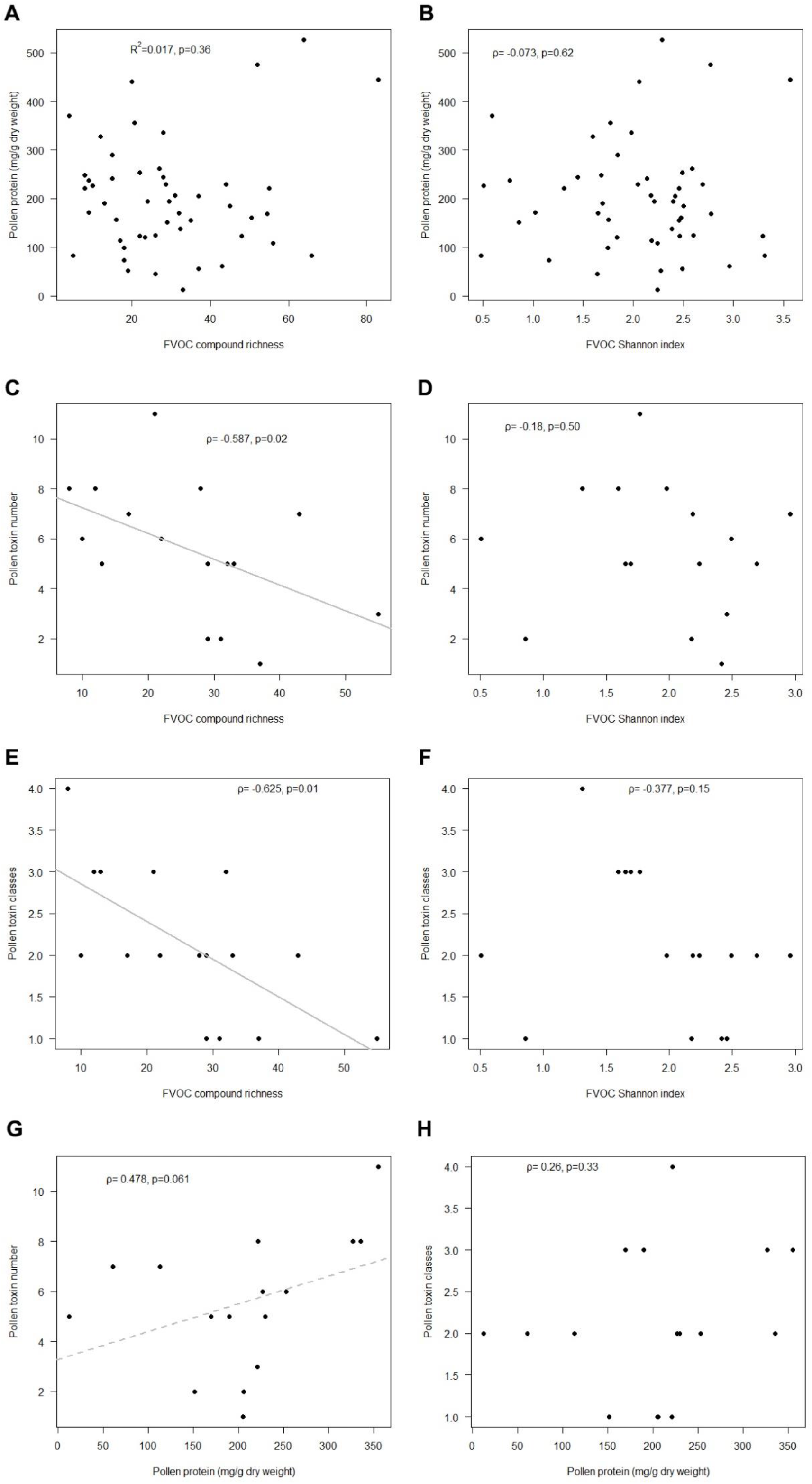
Correlations between FVOC chemodiversity, pollen protein amount and number of described pollen toxins. Correlation between A. pollen protein amount (mg protein/g pollen d.w.) and compound richness (*n* = 49), B. pollen protein amount (mg protein/g pollen d.w.) and FVOC Shannon index (*n* = 49), C. number of pollen toxins and FVOC compound richness (*n* = 16), D. number of pollen toxins and FVOC Shannon index (*n* = 16), E. number of pollen toxin classes and compound richness (*n* = 16), F. number of pollen toxin classes and FVOC Shannon index (*n* = 16), G. number of pollen toxins and pollen protein amount (mg protein/g pollen d.w.) (*n* = 16), H. number of pollen toxin classes and pollen protein amount (mg protein/g pollen d.w.) (*n* = 16). Data was analysed by correlation tests, with *R* (Pearson’s correlation coefficient), *ρ* (Spearman’s rank correlation coefficient) or *p* values given.

In our meta-analysis 2, the pollen protein concentration tended to be higher, when the number of pollen toxins was higher (marginally not significant, Spearman correlation, *ρ* = 0.468, *p* = 0.06, *n* = 16; Fig. 1G). However, no significant relation was found between the protein concentration and the number of toxin classes in pollen (Spearman correlation, *ρ*= 0.26, *p* = 0.33; Fig. 1H). A high number of potential toxins in pollen may be beneficial to defend nutrient-rich pollen more effectively. Losses of pollen due to herbivory would be costly for the plants. These findings thus support the optimal defence hypothesis (McCall and Irwin, 2006, McCall and Fordyce, 2010) in pollen, i.e. nutrient-rich and hence valuable pollen is more defended than nutrient-poor pollen. Testing other hypotheses for plant chemical defences in flowers would require the generation of additional and more comprehensive data from flowers and visitor behaviour (McCall and Irwin, 2006). Among different floral parts, more data can be found for pollen quality in terms of nutrient and defence compounds, whereas it may be likewise interesting to study such traits in nectar and sepals and petals.

More comprehensive approaches, e.g. regarding the multimodal floral “Gestalt” including FVOC, CO_2_, gustatory and visual cues, as done recently (Peach et al., 2019, Kaczorowski et al., 2012, Raguso and Willis, 2005, Katzenberger et al., 2013), are needed. According to the efficacy backup hypothesis, the availability of multimodal cues is costly and may deliver redundant information if all cues are salient. However, as a repetition of information in another language, multimodal cues can enhance decision making (Kulahci et al., 2008) and are particularly decisive for host finding in situations when one cue blurs due to environmental conditions such as wind (Telles et al., 2017, Lawson et al., 2017). Correlations between olfactory and contact cues may also be expected for other plant organs apart from flowers within plant individuals. Chemodiversity can be calculated at different levels and for different organs or tissues. Its use as a multimodal cue and what information it may provide about host plant quality for different types of interaction partners needs further investigation.

## CONCLUSION

The dynamics and plasticity of floral chemistry and insect chemosensation are expected to play a major role in the evolution of plant-insect relationships (Schiestl, 2015). Our review and meta-analyses summarised the essential findings in the role and complexity of various floral chemical displays, including FVOCs, pollen nutrients and pollen toxins, and the impacts these have on flower visitors such as pollinators and florivores. Although the number of studies that consider both pollinators and florivores is limited, we could highlight that some FVOCs, such as linalool and methyl salicylate, are rather attractive for pollinators but repellent for florivores. Nevertheless, there are more shared FVOCs being attractive than repellent for both pollinators and herbivores, increasing the dilemma for plants regarding whom to attract. Chemodiversity is increasingly considered as an important trait mediating interactions within and among species (Müller and Junker, 2022) and has been found to play an important role in mediating interactions with flower visitors (Eilers et al., 2021). Future research should include chemodiversity of different tissues within flowers to gain a better understanding of metabolic, ecological and evolutionary forces driving chemical flower displays.

## ACKNOWLEDGEMENTS

All authors developed the research questions and study design and CM, EJE and RRJ acquired funding. All authors compiled and shared literature. RS compiled and analysed the data, prepared the figure and tables and wrote the first version of the manuscript. All authors revised and commented on the manuscript. We thank Silvia Eckert and other members of the DFG research unit FOR3000 for constructive discussions and valuable comments.

## REFERENCES

Adler LS, Karban R, Strauss SY. 2001. Direct and indirect effects of alkaloids on plant fitness via herbivory and pollination. Ecology, 82: 2032–2044.

Aizen MA, Harder LD. 2007. Expanding the limits of the pollen-limitation concept: effects of pollen quantity and quality. Ecology, 88: 271–281.

Allen B, Kon M, Bar-Yam Y. 2009. A new phylogenetic diversity measure generalizing the Shannon index and its application to phyllostomid bats. The American Naturalist, 174: 236–243.

Annoscia D, Zanni V, Galbraith D, Quirici A, Grozinger C, Bortolomeazzi R, Nazzi F. 2017. Elucidating the mechanisms underlying the beneficial health effects of dietary pollen on honey bees (Apis mellifera) infested by Varroa mite ectoparasites. Scientific Reports, 7: 271–281.

Austel N, Böttcher C, Meiners T. 2021. Chemical defence in Brassicaceae against pollen beetles revealed by metabolomics and flower bud manipulation approaches. Plant, Cell & Environment, 44: 519–534.

Ayestaran A, Giurfa M, de Brito Sanchez MG. 2010. Toxic but drank: gustatory aversive compounds induce post-ingestional malaise in harnessed honeybees. PLoS ONE, 5: e15000.

Barlow SE, Wright GA, Ma C, Barberis M, Farrell IW, Marr EC, Brankin A, Pavlik BM, Stevenson PC. 2017. Distasteful nectar deters floral robbery. Current Biology, 27: 2552-2558.e3.

Barraud A, Vanderplanck M, Nadarajah S, Michez D. 2020. The impact of pollen quality on the sensitivity of bumblebees to pesticides. Acta Oecologica-International Journal of Ecology, 105: 103552.

Bennett JM, Steets JA, Burns JH, Burkle LA, Vamosi JC, Wolowski M, Arceo-Gomez G, Burd M, Durka W, Ellis AG, Freitas L, Li JM, Rodger JG, Stefan V, Xia J, Knight TM, Ashman TL. 2020. Land use and pollinator dependency drives global patterns of pollen limitation in the Anthropocene. Nature Communications, 11: 3999.

Bestea L, Réjaud A, Sandoz JC, Carcaud J, Giurfa M, de Brito Sanchez MG. 2021. Peripheral taste detection in honey bees: What do taste receptors respond to? European Journal of Neuroscience, 54: 4417–4444.

Bhattacharya S, Baldwin IT. 2012. The post-pollination ethylene burst and the continuation of floral advertisement are harbingers of non-random mate selection in Nicotiana attenuata. Plant Journal, 71: 587–601.

Blüthgen N, Klein AM. 2011. Functional complementarity and specialisation: the role of biodiversity in plant-pollinator interactions. Basic and Applied Ecology, 12: 282–291.

Bonvehi JS, Jorda RE. 1997. Nutrient composition and microbiological quality of honeybee-collected pollen in Spain. Journal of Agricultural and Food Chemistry, 45: 725–732.

Borghi M, Fernie AR, Schiestl FP, Bouwmeester HJ. 2017. The sexual advantage of looking, smelling, and tasting good: the metabolic network that produces signals for pollinators. Trends in Plant Science, 22: 338–350.

Bruce TJA, Pickett JA. 2011. Perception of plant volatile blends by herbivorous insects - finding the right mix. Phytochemistry, 72: 1605–1611.

Bukovinszky T, Rikken I, Evers S, Wäckers FL, Biesmeijer JC, Prins HHT, Kleijn D. 2017. Effects of pollen species composition on the foraging behaviour and offspring performance of the mason bee Osmia bicornis (L.). Basic and Applied Ecology, 18: 21–30.

Burdon RCF, Raguso RA, Gegear RJ, Pierce EC, Kessler A, Parachnowitsch AL. 2020. Scented nectar and the challenge of measuring honest signals in pollination. Journal of Ecology, 108: 2132–2144.

Conchou L, Lucas P, Meslin C, Proffit M, Staudt M, Ronou M. 2019. Insect odorscapes: from plant volatiles to natural olfactory scenes. Frontiers in Physiology, 10: 972.

Cook SM, Awmack CS, Murray DA, Williams IH. 2003. Are honey bees’ foraging preferences affected by pollen amino acid composition? Ecological Entomology, 28: 622–627.

Crone MK, Grozinger CM. 2021. Pollen protein and lipid content influence resilience to insecticides in honey bees (Apis mellifera). Journal of Experimental Biology, 224: jeb242040.

Cullen NP, Fetters AM, Ashman TL. 2021. Integrating microbes into pollination. Current Opinion in Insect Science, 44: 48–54.

de Brito Sanchez MG, Giurfa M, Mota TRD, Gauthier M. 2005. Electrophysiological and behavioural characterization of gustatory responses to antennal ‘bitter’ taste in honeybees. European Journal of Neuroscience, 22: 3161–3170.

de Brito Sanchez MG, Ortigao-Farias JR, Gauthier M, Liu FL, Giurfa M. 2007. Taste perception in honeybees: just a taste of honey? Arthropod-Plant Interactions, 1: 69–76.

de Mesquita LX, Maracajá PB, Sakamoto SM, Soto-Blanco B. 2010. Toxic evaluation in honey bees (Apis mellifera) of pollen from selected plants from the semi-arid region of Brazil. Journal of Apicultural Research, 49: 265–269.

DeGrandi-Hoffman G, Corby-Harris V, Carroll M, Toth AL, Gage S, Watkins deJong E, Graham H, Chambers M, Meador C, Obernesser B. 2021. The importance of time and place: nutrient composition and utilization of seasonal pollens by European honey bees (Apis mellifera L.). Insects, 12: 235.

Dobson HEM, Bergström G. 2000. The ecology and evolution of pollen odors. Plant Systematics and Evolution, 222: 63–87.

Dolezal AG, Toth AL. 2018. Feedbacks between nutrition and disease in honey bee health. Current Opinion in Insect Science, 26: 114–119.

Dötterl S, Jürgens A, Seifert K, Laube T, Weissbecker B, Schutz S. 2006. Nursery pollination by a moth in Silene latifolia: the role of odours in eliciting antennal and behavioural responses. New Phytologist, 169: 707–718.

Dudareva N, Klempien A, Muhlemann JK, Kaplan I. 2013. Biosynthesis, function and metabolic engineering of plant volatile organic compounds. New Phytologist, 198: 16–32.

Eckhardt M, Haider M, Dorn S, Müller A. 2014. Pollen mixing in pollen generalist solitary bees: a possible strategy to complement or mitigate unfavourable pollen properties? Journal of Animal Ecology, 83: 588–597.

Eilers EJ, Kleine S, Eckert S, Waldherr S, Müller C. 2021. Flower production, headspace volatiles, pollen nutrients, and florivory in Tanacetum vulgare chemotypes. Frontiers in Plant Science, 11: 611877.

Essenberg CJ. 2021. Intraspecific relationships between floral signals and rewards with implications for plant fitness. AoB PLANTS, 13: plab006.

Farré-Armengol G, Fernandez-Martinez M, Filella I, Junker RR, Peñuelas J. 2020. Deciphering the biotic and climatic factors that influence floral scents: a systematic review of floral volatile emissions. Frontiers in Plant Science, 11: 1154.

Farré-Armengol G, Filella I, Llusia J, Peñuelas J. 2013. Floral volatile organic compounds: between attraction and deterrence of visitors under global change. Perspectives in Plant Ecology, Evolution and Systematics, 15: 56–67.

Farré-Armengol G, Filella I, Llusia J, Peñuelas J. 2016. Bidirectional interaction between phyllospheric microbiotas and plant volatile emissions. Trends in Plant Science, 21: 854–860.

Farré-Armengol G, Filella I, Llusià J, Peñuelas J. 2015. Pollination mode determines floral scent. Biochemical Systematics and Ecology, 61: 44–53.

Fatrcová-Šramková K, Nôžková J, Máriássyová M, Kačániová M. 2016. Biologically active antimicrobial and antioxidant substances in the Helianthus annuus L. bee pollen. Journal of Environmental Science and Health Part B-Pesticides Food Contaminants and Agricultural Wastes, 51: 176–181.

Francis JS, Tatarko AR, Richman SK, Vaudo AD, Leonard AS. 2021. Microbes and pollinator behavior in the floral marketplace. Current Opinion in Insect Science, 44: 16–22.

Giacomini JJ, Connon SJ, Marulanda D, Adler LS, Irwin RE. 2021. The costs and benefits of sunflower pollen diet on bumble bee colony disease and health. Ecosphere, 12: e03663.

Grabe V, Sachse S. 2018. Fundamental principles of the olfactory code. Biosystems, 164: 94–101.

Guiraud M, Hotier L, Giurfa M, de Brito Sanchez MG. 2018. Aversive gustatory learning and perception in honey bees. Scientific Reports, 8: 1343.

Haber AI, Sims JW, Mescher MC, De Moraes CM, Carr DE. 2021. A sensory bias overrides learned preferences of bumblebees for honest signals in Mimulus guttatus. Proceedings of the Royal Society B: Biological Sciences, 288: 20210161.

Hao H, Sun J Fau - Dai J, Dai J. 2013. Dose-dependent behavioral response of the mosquito Aedes albopictus to floral odorous compounds. Journal of Insect Science, 13: 127.

Harrewijn P, Minks AK, Mollema C. 1994. Evolution of plant volatile production in insect-plant relationships. Chemoecology, 5: 55–73.

Hauri KC, Glassmire AE, Wetzel WC. 2021. Chemical diversity rather than cultivar diversity predicts natural enemy control of herbivore pests. Ecological Applications, 31: e02289.

Helletsgruber C, Dötterl S, Ruprecht U, Junker RR. 2017. Epiphytic bacteria alter floral scent emissions. Journal of Chemical Ecology, 43: 1073–1077.

Henselek Y, Eilers EJ, Kremen C, Hendrix SD, Klein A-M. 2018. Pollination requirements of almond (Prunus dulcis): combining laboratory and field experiments. Journal of Economic Entomology, 111: 1006–1013.

Hollister B, Mullin CA. 1998. Behavioral and electrophysiological dose–response relationships in adult western corn rootworm (Diabrotica virgifera virgifera LeConte) for host pollen amino acids. Journal of Insect Physiology, 44: 463–470.

Irwin RE, Adler LS. 2006. Correlations among traits associated with herbivore resistance and pollination: implications for pollination and nectar robbing in a distylous plant. American Journal of Botany, v. 93: 1.

Irwin RE, Cook D, Richardson LL, Manson JS, Gardner DR. 2014. Secondary compounds in floral rewards of toxic rangeland plants: impacts on pollinators. Journal of Agricultural and Food Chemistry, 62: 7335–7344.

Jung JW, Park KW, Ahn YJ, Kwon HW. 2015. Functional characterization of sugar receptors in the western honeybee, Apis mellifera. Journal of Asia-Pacific Entomology, 18: 19–26.

Junker R, Parachnowitsch A. 2015. Working towards a holistic view on flower traits-how floral scents mediate plant-animal interactions in concert with other floral characters. Journal of the Indian Institute of Science, 95: 43–68.

Junker RR. 2016. Multifunctional and diverse floral scents mediate biotic interactions embedded in communities. In: Blande JD, Glinwood RT, eds. Deciphering chemical language of plant communication. Heidelberg: Springer.

Junker RR, Blüthgen N. 2010. Floral scents repel facultative flower visitors, but attract obligate ones. Annals of Botany, 105: 777–782.

Junker RR, Gershenzon J, Unsicker SB. 2011. Floral odor bouquet loses its ant repellent properties after inhibition of terpene biosynthesis. Journal of Chemical Ecology, 37: 1323–31.

Junker RR, Heidinger IMM, Blüthgen N. 2010a. Floral scent terpenoids deter the facultative florivore Metrioptera bicolor (Ensifera, Tettigoniidae, Decticinae). Journal of Orthoptera Research, 19: 69–74.

Junker RR, Höcherl N, Blüthgen N. 2010b. Responses to olfactory signals reflect network structure of flower-visitor interactions. Journal of Animal Ecology, 79: 818–823.

Kaczorowski RL, Leonard AS, Dornhaus A, Papaj DR. 2012. Floral signal complexity as a possible adaptation to environmental variability: a test using nectar-foraging bumblebees, Bombus impatiens. Animal Behaviour, 83: 905–913.

Kantsa A, Raguso RA, Lekkas T, Kalantzi OI, Petanidou T. 2019. Floral volatiles and visitors: a meta-network of associations in a natural community. Journal of Ecology, 107: 2574–2586.

Kárpáti Z, Knaden M, Reinecke A, Hansson BS. 2013. Intraspecific combinations of flower and leaf volatiles act together in attracting hawkmoth pollinators. PLoS ONE, 8: e72805.

Katzenberger TD, Lunau K, Junker RR. 2013. Salience of multimodal flower cues manipulates initial responses and facilitates learning performance of bumblebees. Behavioral Ecology and Sociobiology, 67: 1587–1599.

Kessler D, Diezel C, Clark DG, Colquhoun TA, Baldwin IT. 2013. Petunia flowers solve the defence/apparency dilemma of pollinator attraction by deploying complex floral blends. Ecology Letters, 16: 299–306.

Kessler SC, Tiedeken EJ, Simcock KL, Derveau S, Mitchell J, Softley S, Stout JC, Wright GA. 2015. Bees prefer foods containing neonicotinoid pesticides. Nature, 521: 74–U145.

Khan KA, Ghramh HA, Ahmad Z, El-Niweiri MAA, Mohammed MEA. 2021. Honey bee (Apis mellifera) preference towards micronutrients and their impact on bee colonies. Saudi Journal of Biological Sciences, 28: 3362–3366.

Kim JH, Mullin CA. 2007. An isorhamnetin rhamnoglycoside serves as a costimulant for sugars and amino acids in feeding responses of adult western corn rootworms (Diabrotica virgifera virgifera) to corn (Zea mays) pollen. Journal of Chemical Ecology, 33: 501–512.

Knauer AC, Kokko H, Schiestl FP. 2021. Pollinator behaviour and resource limitation maintain honest floral signalling. Functional Ecology, 35: 2536–2549.

Knauer AC, Schiestl FP. 2015. Bees use honest floral signals as indicators of reward when visiting flowers. Ecology Letters, 18: 135–143.

Knight TM, Steets JA, Vamosi JC, Mazer SJ, Burd M, Campbell DR, Dudash MR, Johnston MO, Mitchell RJ, Ashman TL. 2005. Pollen limitation of plant reproduction: pattern and process. Annual Review of Ecology Evolution and Systematics, 36: 467–497.

Knudsen JT, Eriksson R, Gershenzon J, Stahl B. 2006. Diversity and distribution of floral scent. Botanical Review, 72: 1–120.

Knudsen JT, Gershenzon J. 2006. The chemical diversity of floral scent. In: Pichersky E, Dudareva N, eds. Biology of floral scent. 2nd Edition ed. Boca Raton: CRC Press.

Kriesell L, Hilpert A, Leonhardt SD. 2017. Different but the same: bumblebee species collect pollen of different plant sources but similar amino acid profiles. Apidologie, 48: 102–116.

Kulahci IG, Dornhaus A, Papaj DR. 2008. Multimodal signals enhance decision making in foraging bumble-bees. Proceedings of the Royal Society B-Biological Sciences, 275: 797–802.

Larue A-AC, Raguso RA, Junker RR. 2016. Experimental manipulation of floral scent bouquets restructures flower–visitor interactions in the field. Journal of Animal Ecology, 85: 396–408.

Lawson DA, Whitney HM, Rands SA. 2017. Colour as a backup for scent in the presence of olfactory noise: testing the efficacy backup hypothesis using bumblebees (Bombus terrestris). Royal Society Open Science, 4: 170996.

Liolios V, Tananaki C, Dimou M, Kanelis D, Goras G, Karazafiris E, Thrasyvoulou A. 2015. Ranking pollen from bee plants according to their protein contribution to honey bees. Journal of Apicultural Research, 54: 582–592.

Lucas-Barbosa D. 2016. Integrating studies on plant-pollinator and plant-herbivore interactions. Trends in Plant Science, 21: 125–133.

Lundgren JG, Wiedenmann RN. 2004. Nutritional suitability of corn pollen for the predator Coleomegilla maculata (Coleoptera: Coccinellidae). Journal of Insect Physiology, 50: 567–575.

Manson JS, Otterstatter MC, Thomson JD. 2010. Consumption of a nectar alkaloid reduces pathogen load in bumble bees. Oecologia, 162: 81–89.

May CE, Dus M. 2021. Confection confusion: interplay between diet, taste, and nutrition. Trends in Endocrinology and Metabolism, 32: 95–105.

McCall AC, Fordyce JA. 2010. Can optimal defence theory be used to predict the distribution of plant chemical defences? Journal of Ecology, 98: 985–992.

McCall AC, Irwin RE. 2006. Florivory: the intersection of pollination and herbivory. Ecology Letters, 9: 1351–1365.

Mitchell BK, Itagaki H, Rivet MP. 1999. Peripheral and central structures involved in insect gustation. Microscopy Research and Technique, 47: 401–415.

Müller C, Junker RR. 2022. Chemical phenotype as important and dynamic niche dimension of plants. New Phytologist, 234: 1168–1174.

Muth F, Breslow PR, Masek P, Leonard AS. 2018. A pollen fatty acid enhances learning and survival in bumblebees. Behavioral Ecology, 29: 1371–1379.

Muth F, Francis JS, Leonard AS. 2016. Bees use the taste of pollen to determine which flowers to visit. Biology Letters, 12: 20160356.

Nicolson SW. 2011. Bee food: the chemistry and nutritional value of nectar, pollen and mixtures of the two. African Zoology, 46: 197–204.

Nicolson SW, Human H. 2013. Chemical composition of the ‘low quality’ pollen of sunflower (Helianthus annuus, Asteraceae). Apidologie, 44: 144–152.

Palmer-Young EC, Farrell IW, Adler LS, Milano NJ, Egan PA, Junker RR, Irwin RE, Stevenson PC. 2019. Chemistry of floral rewards: intra-and interspecific variability of nectar and pollen secondary metabolites across taxa. Ecological Monographs, 89: 1–20.

Peach DAH, Gries R, Zhai HM, Young N, Gries G. 2019. Multimodal floral cues guide mosquitoes to tansy inflorescences. Scientific Reports, 9: 3908.

Petrén H, Köllner TG, Junker RR. 2023. Quantifying chemodiversity considering biochemical and structural properties of compounds with the R package chemodiv. New Phytologist, in press. doi:10.1101/2021.09.30.462509.

Pichersky E, Noel JP, Dudareva N. 2006. Biosynthesis of plant volatiles: nature’s diversity and ingenuity. Science, 311: 808–811.

Poivet E, Gallot A, Montagne N, Senin P, Monsempes C, Legeai F, Jacquin-Joly E. 2021. Transcriptome profiling of starvation in the peripheral chemosensory organs of the crop pest Spodoptera littoralis caterpillars. Insects, 12: 573.

Proffit M, Bessiere JM, Schatz B, Hossaert-McKey M. 2018. Can fine-scale post-pollination variation of fig volatile compounds explain some steps of the temporal succession of fig wasps associated with Ficus racemosa? Acta Oecologica-International Journal of Ecology, 90: 81–90.

Quinlan G, Milbrath M, Otto C, Smart A, Iwanowicz D, Cornman RS, Isaacs R. 2021. Honey bee foraged pollen reveals temporal changes in pollen protein content and changes in forager choice for abundant versus high protein flowers. Agriculture Ecosystems & Environment, 322: 107645.

Raguso RA. 2008a. Start making scents: the challenge of integrating chemistry into pollination ecology. Entomologia Experimentalis Et Applicata, 128: 196–207.

Raguso RA. 2008b. Wake up and smell the roses: the ecology and evolution of floral scent. Annual Review of Ecology Evolution and Systematics, 39: 549–569.

Raguso RA, Willis MA. 2005. Synergy between visual and olfactory cues in nectar feeding by wild hawkmoths, Manduca sexta. Animal Behaviour, 69: 407–418.

Reinhard J, Srinivasan MV, Zhang SW. 2004. Olfaction: scent-triggered navigation in honeybees. Nature, 427: 411–411.

Rivest S, Forrest JRK. 2020. Defence compounds in pollen: why do they occur and how do they affect the ecology and evolution of bees? New Phytologist, 225: 1053–1064.

Roulston TH, Buchmann SL. 2000. A phylogenetic reconsideration of the pollen starch-pollination correlation. Evolutionary Ecology Research, 2: 627–643.

Roulston TH, Cane JH, Buchmann SL. 2000. What governs protein content of pollen: pollinator preferences, pollen-pistil interactions, or phylogeny? Ecological Monographs, 70: 617–643.

Ruedenauer FA, Biewer NW, Nebauer CA, Scheiner M, Spaethe J, Leonhardt SD. 2021. Honey bees can taste amino and fatty acids in pollen, but not sterols. Frontiers in Ecology and Evolution, 9: 684175.

Ruedenauer FA, Raubenheimer D, Kessner-Beierlein D, Grund-Mueller N, Noack L, Spaethe J, Leonhardt SD. 2020. Best be(e) on low fat: linking nutrient perception, regulation and fitness. Ecology Letters, 23: 545–554.

Ruedenauer FA, Spaethe J, van der Kooi CJ, Leonhardt SD. 2019. Pollinator or pedigree: which factors determine the evolution of pollen nutrients? Oecologia, 191: 349–358.

Salzmann C, xa, Brown A, Schiestl F, xa. 2006. Floral scent emission and pollination syndromes: evolutionary changes from food to sexual deception. International Journal of Plant Sciences, 167: 1197–1204.

Sasidharan A, Venkatesan R. 2020. Olfactory cues as functional traits in plant reproduction. In: Tandon R, Shivanna KR, Koul M, eds. Reproductive ecology of flowering plants: patterns and processes. Singapore: Springer Nature.

Schiestl FP. 2010. The evolution of floral scent and insect chemical communication. Ecology Letters, 13: 643–656.

Schiestl FP. 2015. Ecology and evolution of floral volatile-mediated information transfer in plants. New Phytologist, 206: 571–577.

Schiestl FP, Ayasse M. 2001. Post-pollination emission of a repellent compound in a sexually deceptive orchid: a new mechanism for maximising reproductive success? Oecologia, 126: 531–534.

Schiestl FP, Johnson SD. 2013. Pollinator-mediated evolution of floral signals. Trends in Ecology & Evolution, 28: 307–315.

Schmehl DR, Teal PEA, Frazier JL, Grozinger CM. 2014. Genomic analysis of the interaction between pesticide exposure and nutrition in honey bees (Apis mellifera). Journal of Insect Physiology, 71: 177–190.

Schmidt HR, Benton R. 2020. Molecular mechanisms of olfactory detection in insects: beyond receptors. Open Biology, 10: 200252.

Sokolinskaya EL, Kolesov DV, Lukyanov KA, Bogdanov AM. 2020. Molecular principles of insect chemoreception. Acta Naturae, 12: 81–91.

Spear DM, Silverman S, Forrest JRK. 2016. Asteraceae pollen provisions protect Osmia mason bees (Hymenoptera: Megachilidae) from brood parasitism. American Naturalist, 187: 797–803.

Stevenson PC. 2020. For antagonists and mutualists: the paradox of insect toxic secondary metabolites in nectar and pollen. Phytochemistry Reviews, 19: 603–614.

Strauss SY, Irwin RE, Lambrix VM. 2004. Optimal defence theory and flower petal colour predict variation in the secondary chemistry of wild radish. Journal of Ecology, 92: 132–141.

Strauss SY, Whittall JB. 2010. Non-pollinator agents of selection on floral traits. Ecology and Evolution of Flowers: 120–138.

Telles FJ, Corcobado G, Trillo A, Rodriguez-Gironés MA. 2017. Multimodal cues provide redundant information for bumblebees when the stimulus is visually salient, but facilitate red target detection in a naturalistic background. PloS ONE, 12: e0184760.

Terry I, Walter GH, Moore C, Roemer R, Hull C. 2007. Odor-mediated push-pull pollination in cycads. Science, 318: 70.

Theis N, Adler LS. 2012. Advertising to the enemy: enhanced floral fragrance increases beetle attraction and reduces plant reproduction. Ecology, 93: 430–435.

Theis N, Lerdau M, Raguso RA. 2007. The challenge of attracting pollinators while evading floral herbivores: patterns of fragrance emission in Cirsium arvense and Cirsium repandum (Asteraceae). International Journal of Plant Sciences, 168: 587–601.

Tosi S, Nieh JC, Sgolastra F, Cabbri R, Medrzycki P. 2017. Neonicotinoid pesticides and nutritional stress synergistically reduce survival in honey bees. Proceedings of the Royal Society B-Biological Sciences, 284: 20171711.

Twaij BM, Hasan MN. 2022. Bioactive secondary metabolites from plant sources: types, synthesis, and their therapeutic uses. International Journal of Plant Biology, 13: 4–14.

Underwood N, Hambäck PA, Inouye BD. 2020. Pollinators, herbivores, and plant neighborhood effects. The Quarterly Review of Biology, 95: 37 –57.

Vaudo AD, Patch HM, Mortensen DA, Tooker JF, Grozinger CM. 2016. Macronutrient ratios in pollen shape bumble bee (Bombus impatiens) foraging strategies and floral preferences. Proceedings of the National Academy of Sciences of the United States of America, 113: E4035–E4042.

Vaudo AD, Tooker JF, Patch HM, Biddinger DJ, Coccia M, Crone MK, Fiely M, Francis JS, Hines HM, Hodges M, Jackson SW, Michez D, Mu JP, Russo L, Safari M, Treanore ED, Vanderplanck M, Yip E, Leonard AS, Grozinger CM. 2020. Pollen protein: lipid macronutrient ratios may guide broad patterns of bee species floral preferences. Insects, 11: 132.

Wetzel WC, Whitehead SR. 2020. The many dimensions of phytochemical diversity: linking theory to practice. Ecology Letters, 23: 16–32.

Whitehead MR, Peakall R. 2009. Integrating floral scent, pollination ecology and population genetics. Functional Ecology, 23: 863–874.

Wolters-Arts M, Lush WM, Mariani C. 1998. Lipids are required for directional pollen-tube growth. Nature, 392: 818–821.

Wright GA, Mustard JA, Simcock NK, Ross-Taylor AA, McNicholas LD, Popescu A, Marion-Poll F. 2010. Parallel reinforcement pathways for conditioned food aversions in the honeybee. Current Biology, 20: 2234–40.

Wright GA, Schiestl FP. 2009. The evolution of floral scent: the influence of olfactory learning by insect pollinators on the honest signalling of floral rewards. Functional Ecology, 23: 841–851.

Zarchin S, Dag A, Salomon M, Hendriksma HP, Shafir S. 2017. Honey bees dance faster for pollen that complements colony essential fatty acid deficiency. Behavioral Ecology and Sociobiology, 71: 172.

Zu P, García-García R, Schuman M, Saavedra S, Melián CJ. 2022. Plant-insect chemical communication in ecological communities: an information theory perspective. Journal of Systematics and Evolution, in press. doi:10.1101/2021.09.30.462509.

