## Supplement S1 for "Floral volatiles evoke partially similar responses in both florivores and pollinators and are correlated with non-volatile reward chemicals"

**Methods for literature review and meta-analyses**

For our literature review and meta-analyses, data were obtained by searching in the ISI Web of Science (1945–2020), Google Scholar, Google Web Search and the database SCENTbase, University of Gothenburg, http://www2.botany.gu.se/SCENTbase.html (Knudsen et al., 2006). The following search terms were used in combinations: "flower", "floral", "scent", "odour", "volatile", "VOC", "headspace", "pollinator", "florivore", "floral herbivore", "GC-EAD"," EAG", "electrophysiology", "olfactometry", "choice behaviour", "poll* protein", "poll* nutri*", "poll* tox*".

For our meta-analysis 1, we addressed the question of whether mutualistic (i.e., pollinators) and antagonistic (i.e., florivores) flower-visiting insects differ in their ability to detect and show behavioural responses towards floral volatile organic compounds (FVOCs) of the same plant genera. Pollinators were defined as such if it was reported that they cross-transfer pollen within the same plant species, while florivores were defined as such if they were reported to damage floral tissues (e.g., sepals, petals, unripe seeds), or if they were found to act as nectar or pollen thieves. Accordingly, we linked qualitative FVOC data to electrophysiological and behavioural responses of pollinators and florivores towards these compounds within the respective plant species and genera. Data were selected only for those plant species, for which neuro-ethological data covering both pollinators and florivores was available. The required combination of data for these meta-analyses was only available for a small set of plant species ($n$ = 8). Therefore, data from closely related plant species within the same plant genera was also included and data was pooled to the genus level for both meta-analyses.

For meta-analysis 2, we expanded the dataset to include all plant species, for which both FVOC data and data on pollen protein amount and/or stored specialised toxic metabolites of pollen (defined by Rivest and Forrest, 2020) as belonging to certain classes of specialised metabolites and known for a deterrent role in at least one species) were available.

We did not include data in the meta-analyses with insufficient description of the methods, insufficient number of replicates or data from studies which lacked a suitable control for determining a positive or negative behavioural response.

**Pollinator and florivore responses to FVOCs: meta-analysis 1**

Within the literature selection for FVOCs from different plant species, we searched for neuroethological data that were available for both pollinators and florivores. Specifically, the literature was searched for data on pollinator and florivore electrophysiology (i.e., neuro-peripheral compound detection ability) and behaviour towards FVOCs. In total, we found sufficiently extensive datasets for eight plant genera, namely *Brassica* (Brassicaceae), *Cirsium* and *Helianthus* (Asteraceae), Cucurbita (Cucurbitaceae), *Daucus* (Apiaceae), *Dichaea pendula* (Orchidaceae), *Fragaria* (Rosaceae) and *Nicotiana* (Solanaceae). In two of these genera (i.e. *Daucus* and *Dichaea*), no detection data was available for one of either pollinators or florivores, but behavioural data was available for both groups.

Data on detection were assigned to binary values (1 or 0), depending on whether the floral volatile compounds elicited a response (1) or not (0). This data was collected from electrophysiological assays including gas chromatography-electroantennographic detection (GC-EAD) or electroantennography (EAG) and behavioural data (either positive or negative responses indicate a response of 1 for detection). Behavioural data was either based on choice or non-choice preference tests or trap assays. Compounds were assigned either +, - or 0, based on whether they were significantly attractive (+), repellent (-) or provoked no measurable behaviour (0), respectively, in reference to a control.

For the detection data, the percentage of unique, active FVOC tests (test of a unique FVOC against a unique insect species) was calculated for pollinators or florivores for each plant genus among all unique FVOC tests for that insect group. For the behavioural data, the percentage of attractive and repellent FVOC responses was individually calculated, for pollinators or florivores and for each plant genus, among all unique FVOC tests. Several FVOCs were present in more than one plant genus and the results for the same insect species and FVOCs across different plant species were merged only in the total. Therefore, each test (each value of n) represents a unique compound-insect interaction. Some of the studies using electrophysiology did not state the number of tested FVOCs but showed GC chromatograms from which the number of unknown compounds was obtained by counting the peaks. For the electrophysiology data, FVOCs determined in the GC chromatograms that did not provoke measurable EAG spikes were considered as not detectable by the chemosensory organs of the respective insect species. Electrophysiology data of FVOCs that were not identified by the authors were included in our dataset as sequentially numbered unknown compounds. Behavioural data that suggested that an FVOC is also detected (by being attractive or repellent in a behavioural test) was also extrapolated to detection by assigning a value of (1) for the detection of that FVOC.

We found behavioural data for 112 and 159 FVOC-insect tests, for pollinators and florivores, respectively. In this data, only compounds directly tested against the corresponding pollinators or florivores were considered as tested compounds. We did not find studies on behavioural responses of florivores towards *Cucurbita* spp. and *Helianthus* spp. FVOCs. All data were treated uniformly between pollinators and florivores to ensure that data evaluation was not biased between the two groups.

**Relationships between FVOCs and pollen chemicals: meta-analysis 2**

In meta-analysis 2, we tested compound richness (i.e. number) or compound diversity of FVOCs, stored toxins in pollen, and pollen protein for correlations ($n$ = 16 plants had toxin data available, $n$ = 49 plants had data for pollen protein and FVOCs). Plant species were selected based on the joint availability of data on their FVOC profile (as a measure of chemodiversity), their pollen protein content (as a measure of reward quality, Pamminger et al., 2019) and stored toxin number). As a measure for chemodiversity of FVOCs, we used the compound richness and Shannon diversity, calculated based on the FVOC data from various sources (using the composition data of the FVOCs from GC-MS chromatograms). The Shannon diversity index was calculated from the FVOC data according to the formula

$$H_s = -\Sigma\, p_i * \ln p_i,$$

where $p$ is the relative abundance of an FVOC i (Shannon and Weaver, 1949), using the vegan package in R Studio (see below). When multiple sources were available for FVOCs or pollen protein belonging to the same plant species, the FVOC diversity index (compound richness or Shannon index) or pollen protein was averaged for that species across the different sources. Pollen toxins were grouped into their respective chemical classes (e.g., alkaloids, terpenoids etc.) and the numbers of pollen toxins and toxin classes were tested for correlations with FVOC richness, Shannon diversity and pollen protein amount.

### STATISTICAL ANALYSES

All analyses were performed using R v4.0.3 (https://www.r-project.org) and R Studio v1.4 (https://posit.co). Pearson's chi-square test was used to identify the relation between insect group, i.e., pollinator versus florivore, and the detection (EAD) or attractiveness vs. repellence (behaviour) of floral volatiles from the plant genera or the total from the different plant species or the different FVOCs. Binomial tests were used to test the distribution of shared attractive and repellent FVOCs between pollinators and florivores, as counts of shared active compounds could be divided between shared attractive and shared repellent, and are hence independent.

For correlation analyses, data was first checked for a normal distribution using the Shapiro-Wilk test (package stats) and Levene's test for homogeneity of variances (package car). Data that gave a normal distribution based on these tests were assumed to be parametric. Accordingly, Pearson's or Spearman's rank correlation analyses (package stats) were performed to calculate the correlation of the relation between the various reward traits, including pollen protein and stored defences and the FVOC diversity indices of the plant species.

### REFERENCES


**Knudsen JT, Eriksson R, Gershenzon J, Stahl B. 2006**. Diversity and distribution of floral scent. *Botanical Review,* **72**: 1-120.

**Pamminger T, Becker R, Himmelreich S, Schneider CW, Bergtold M. 2019**. Pollen report: quantitative review of pollen crude protein concentrations offered by bee pollinated flowers in agricultural and non-agricultural landscapes. *PeerJ,* **7**: e7394.

**Rivest S, Forrest JRK. 2020**. Defence compounds in pollen: why do they occur and how do they affect the ecology and evolution of bees? *New Phytologist,* **225**: 1053-1064.

**Shannon CE, Weaver W. 1949**. *The mathematical theory of communication.* Champaign: University of Illinois Press.
